# Serum proteomics expands on high-affinity antibodies in immunized rabbits than deep B-cell repertoire sequencing alone

**DOI:** 10.1101/833871

**Authors:** Stefano R. Bonissone, Thiago Lima, Katherine Harris, Laura Davison, Brian Avanzino, Nathan Trinklein, Natalie Castellana, Anand Patel

**Affiliations:** Digital Proteomics LLC, San Diego, CA, USA; Teneobio Inc., Menlo Park, CA, USA

## Abstract

Rabbits are a model for immunology studies, and monoclonal antibodies developed from rabbits have been sought after to empower immunoassays in a variety of applications. High-throughput characterization of circulating serum antibodies in response to specific antigens is highly impactful for both humoral immunology studies and antibody development. A combination of high throughput sequencing of antibody transcripts from B cells and proteomic analysis of serum antibodies, an approach referred to as immunoproteogenomics, is applied to profile the immune response of rabbits to *β*-galactosidase (Beta-gal) in both recombinant antigen and peptide antigen immunization formats. The use of intact protein antigen resulted in observing 56.3% more heavy chains CDR3s in serum than immunization with peptide antigens. Additionally, sampling peripheral blood mononuclear cells (PBMCs) for B-cell repertoire sequencing at different time points throughout the immunization was found to capture 47.8%-72.8% of total proteomically observed heavy chain CDR3s, and would serve well in replacing sequencing the B cell rich, but more difficult to access spleen or bone marrow compartments. Despite B-cell repertoire sequencing to depths of 2M to 10M reads, we found proteomic evidence supporting at least 10% of serum antibodies are still missed. Further improvements to proteomic analysis techniques would enable more precise characterization of antibodies circulating in serum and determine antibody protein sequences missed by repertoire sequencing.

## 1 Introduction

High-throughput sequencing of antibody transcripts originating from the rearranged immunoglobulin (Ig) locus in B-cell populations (Rep-seq) has uncovered vastly diverse antibody sequences. Rep-seq has shed light on mechanisms of immunity ranging from understanding quiescent stasis [16], activation in response to vaccines [15, 21], differences in vaccine response of identical twins [37], response to chronic HIV infection [41] and immunization [14], engagement within the tumor microenvironment [5], and breaking of self-tolerance for autoimmune diseases [35, 7]. However, the Ig transcripts from cells sampled from various compartments and tissues are only a small component of the humoral immune response. Antibodies circulating in the serum more accurately reflect the state of the humoral immune response. Emil von Behring and Kitasato Shibasaburo were the first to report the presence of diptheria and tentanus neutralizing substances extracted from animal blood in the 1890s [3]. To date, few groups have analyzed circulating antibodies in the serological compartment in high-throughput and at sequence-level resolution.

Tandem mass spectrometry (LC-MS/MS) is currently the only high-throughput technology capable of characterizing protein sequences. Direct protein sequencing of monoclonal antibodies (mAbs) [1, 6, 36] can be achieved by combining standard bottom-up proteomic approaches with computational analysis. However, polyclonal antibodies (pAbs) found in a natural immune response are highly complex and few attempts to directly sequence them have succeeded [13]. Immunoproteogenomics is an approach that combines transcriptomics with proteomics to determine the repertoire of antibodies present in sample.To our knowledge, Obermeier et al. 2008 [29] was the first study to combine transcript sequences from B cells in the cerebral spinal fluid (CSF) and match them against mass spectra from IgGs purified from CSF as well as serum. In patients diagnosed with multiple sclerosis, the authors found strong overlap between oligoclonal serum antibodies and Ig transcripts from B cells. In a later immunoproteogenomics study, Cheung et al., 2012 [8] identified serum antibodies from hyperimmunized rabbits and mice. The same group showed the potential of immunoproteogenomics in human by identifying functional serum-derived antibodies against CMV and influenza [32]. The Georgiou lab further developed immunoproteogenomics to demonstrate identification of serum antibodies in immunized rabbit [39] and human vaccinations [23]. Similar to CSF, affinity-purified antibodies from serum had strong overlap with Ig transcripts from B cells collected from bone marrow and peripheral blood mononuclear cells (PBMCs). Lavinder et al., argued that the LC-MS/MS limit of detection is at the 0.1nM theoretical ceiling for neutralization affinity of antibodies, and their immunoproteogenomic approach sufficiently captures the relevant antibody clonotypes in the serological compartment [23]. Additional groups have applied immunoproteogenomics to llama [11], CMV infected human [31], and human pemphigus [7].

In this study, we apply a immunoproteogenomic approach to systematically characterize the immune response of rabbits to two types of immunogens: peptides conjugated to an immunogenic carrier protein and an intact protein. We compare the antibody proteomic measurements to the B-cell repertoire derived from next-generation sequencing (Rep-seq). Rabbits are well suited for immunoproteogenomic experiments, due to the larger number of B cells and quantity of serum antibodies that can be collected from a single individual. Additionally, rabbit antibodies empower a number of research and therapeutic applications. Polyclonal rabbit antibodies are used extensively for immunohistochemistry, western blot, and flow cytometry assays, with most antibody catalog companies pAb products being sourced from rabbits. Companies, such as AbCam, Inc., Cell Signaling Technology, Inc. (>4000 rabbit monoclonal antibodies listed in online catalog), Agilent, and ThermoFisher Scientific have pursued monoclonal rabbit antibody development for highly specific and reproducible assays. In medicine, rabbits have been used to improve HIV-1 vaccine design [2], and humanized rabbit mAbs developed by Apexigen Inc. are undergoing clinical trials to treat cancer and ocular diseases [25]. Interest in rabbit antibodies is due to their reported higher affinities and unique specificity [19, 38]. Although, Landry et al. [20] reported the improvement may only be slight, with rabbit mAbs generated against linear peptides having on 20-200 pM binding (across 1450 rabbit mAbs), compared to mouse mAbs with 30-300pM binding (across 46 mouse mAbs). However, even they noted only rabbit mAbs had binding close to the 1pM detection limit of surface plasmon resonance. Also, a number of head-to-head studies comparing mouse to rabbit monoclonals against the same human antigen for IHC have shown rabbit antibodies have superior sensitivity [38]. There are a number of reasons for affinity differences between antibodies from different immunized hosts, and a contributing factor is differences in B cell lymphogenesis. In young rabbits, the pre-immune antibody repertoire is generated in the bone marrow where progenitor B cells undergo VDJ rearrangement. B cells migrate to gut-associated lymphoid tissue (GALT) where somatic hypermutation and gene conversion shape a diverse primary immune repertoire [26]. Further affinity maturation occurs in spleen and lymph nodes in response to immunizations or infections. The three-step process for developing a diverse repertoire is distinct from other mammals, like mouse and humans, and a likely contributing factor to observing high affinity antibodies in rabbits.

In this investigation, the immune response of rabbits immunized with recombinant *β*-galactosidase (Beta-gal) antigen is tracked by performing Rep-seq on PBMCs collected at multiple time points and compartments. Since high titer antibodies to the antigen are found in serum during immunizations, B cells that generate the antigen-specific antibodies are also expected to be abundant and transcriptionally active in PBMCs, and should be detected by Rep-seq. However, immunoproteogenomics studies have shown high affinity antibodies are not transcriptionally the most abundant [8, 31]. Differing from previous studies, this investigation profiles the rabbit immune response with deeper B-cell repertoire sequencing, and across multiple time points during the immunization and tissues. The data presented will show the benefits and caveats of immunoproteogenomic studies and underscores the importance of accurate data, timing of B cell collections, and sequencing depth for tracking the immune response to a specific antigen. Importantly, Rep-seq studies in humans and large animals such as goats and camelids are typically restricted to only sampling PBMCs. The data will show blood collections from multiple time points during maturation of an immune response provides a suitable proxy for B cells found in spleen and bone marrow. As the immune response to Beta-gal in both intact protein and peptide antigens formats are characterized, the data reveals differences in antigen-specific clonal diversity for each format, and trade-offs for adequately capturing high affinity antibodies observed in blood versus tissue. Lastly, there is compelling proteomic evidence of high affinity antibodies that are not represented by any sampled B cells, despite high depth of B-cell sequencing.

## 2 Materials and Methods

### 2.1 Rabbit subjects and antigen

Data was generated from immunizations of three juvenile New Zealand white rabbits: two rabbits with recombinant *Escherichia coli* Beta-gal (*β*-galactosidase from Sigma Aldrich, Inc.), and one rabbit with two linear peptides RNSEEARTDRPSQQLRSLNGE (peptide A) at positions 38-48 and DVAPQGKQLIELPELPQPES (peptide B) at positions 672-691 of Beta-gal. The three rabbits of random sex were immunized in complete Freund’s adjuvant. Additional boosters of incomplete Freund’s adjuvant were administered at weeks 3, 6, and 10. Serum titers were measured with ELISA for productive IgG response, and rabbits producing highest titer against the antigen at 7 weeks were selected for analysis. Bleeds were performed pre- and post-boost, together with a final bleed at 14 weeks. PBMCs were isolated with Ficoll gradient from pre-immune, pre-boost, and post-boost bleeds. From the final bleed, 15mL was reserved for PBMC isolation, and remaining serum was used for Ig purification. Bone marrow and spleen were harvested at 14 weeks. Repertoire sampling for the two immunized rabbits with the highest antibody titers is shown in Figure 1a.

**Figure 1:**
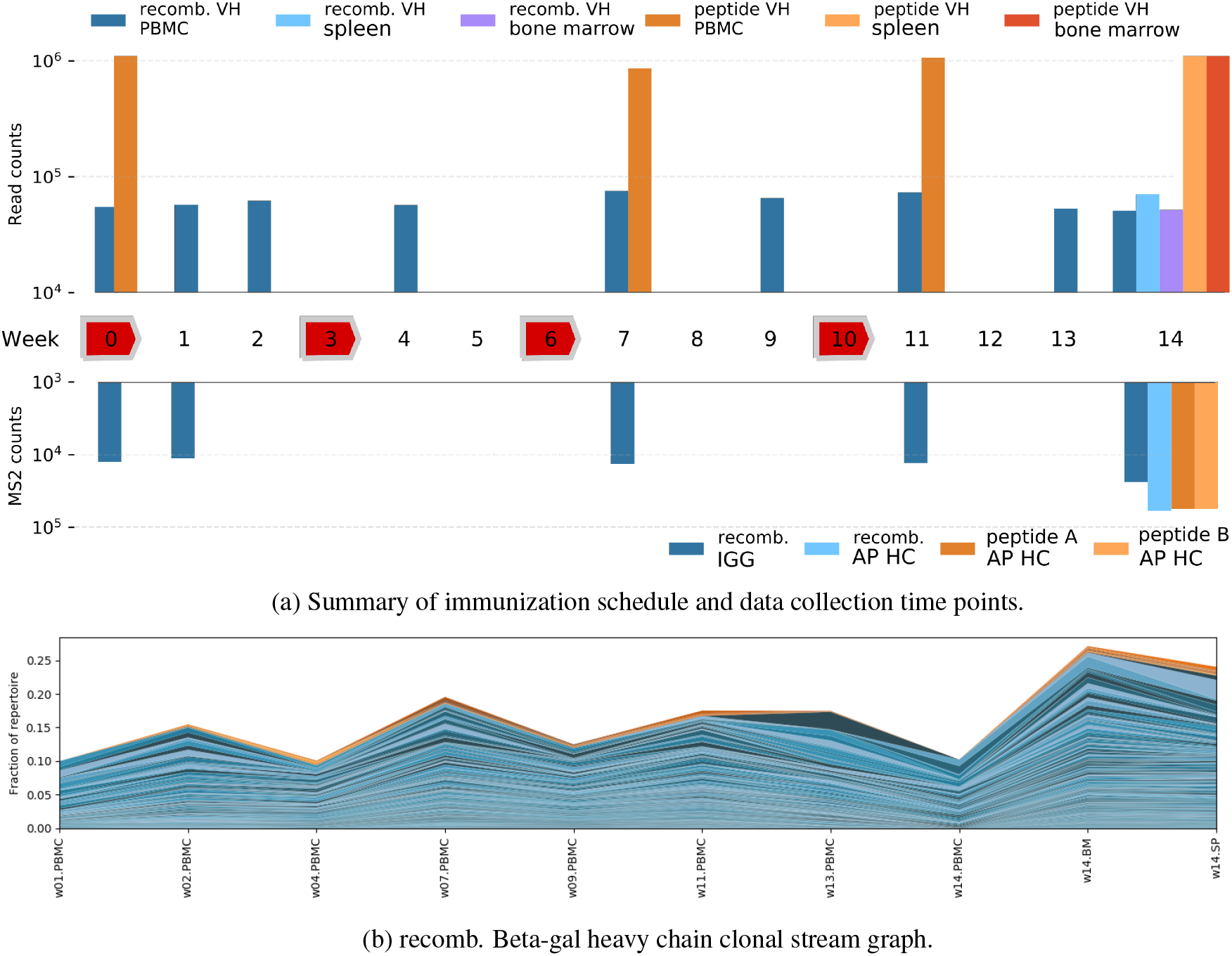
Immunization schedule with time points of Rep-seq and MS/MS data collection. (a) Immunization spans 14 weeks with boosts at 3, 6, and 10 weeks (red). B cells collected at nine time points for recombinant Beta-gal immunization (blue), and four time points for peptide Beta-gal immunization (orange). Serum proteomic samples were collected from five different time points. Only variable heavy transcripts and MS/MS for heavy chain is shown. (b) Stream graph of heavy chain clones that appear in at least two recombinant Beta-gal repertoires where stream height is fraction of sequences in the respective repertoire. Colors represent clones observed only in repertoires (blue), and proteomically observed clones (orange).

### 2.2 Ig transcript sequencing

RNA was extracted from PBMCs, spleen, and bone marrow samples stored in RNALater. RNA were enriched for Ig transcripts by multiplex polymerase chain reactions (PCR) as performed in [40, 19]. Gene-specific primers were designed to complement the Ig leaders and the IgG constant region to ensure full transcript sequencing. The 5’ end of primer sequences contain a sequencing library barcode and Illumina P5 and P7 adaptors to allow for direct paired-end sequencing read (300bp × 300bp) on the Illumina MiSeq sequencer. For each sample, variable heavy chain was amplified in separate PCRs from kappa light chain. The sequencing depth for each library from immunization samples is shown in Table S1.

### 2.3 Proteomic analysis of serum antibodies

Serum antibodies were purified from 25mL of remaining serum of final bleeds using antigen conjugated to NHS-activated agarose resin (ThermoFisher). Columns were washed three times with 1X PBS buffer. Antibody bound to the column was eluted with 20mL of 0.25M glycine HCl pH 1.85 elution buffer into 20 fractions. Fractions were buffer exchanged and concentrated using 5kDa MWCO filter (Corning Spin-X), and quantified by Qubit fluorometer (Life Technologies). A single fraction was selected and ran in multiple lanes on a reducing SDS-PAGE. Gel bands for heavy and light chains were excised and in-gel digested using trypsin (Promega), chymotrypsin (Promega), elastase (Promega), and pepsin (Worthington). Digested peptides were individually subjected to LC-MS/MS with 2hr gradients on an Orbitrap Fusion Lumos Tribid (ThermoFisher). MS/MS was acquired in data-dependent precursor selection (see Table S1 for MS/MS spectral counts per run). In the recombinant Beta-gal immunization, additional serum samples from weeks 0, 1, 7, and 11 from the two rabbits were pooled to start with 200ug for each time point and then, purified with Protein G, treated with IdeS, analyzed by SDS-PAGE, and gel excised to obtain Fab fractions. The fractions were digested with trypsin, chymotrypsin, and elastase and MS/MS data was collected on an Orbitrap Velos in CID mode on a 1hr gradient. From the final bleed at week 14, Fab fractions were obtained from Beta-gal affinity purification of 25mL of pooled serum and Protein G purification of 7.5mL of pooled serum. The pooled final bleeds were digested with seven enzymes, and analyzed on an Orbitrap Velos in CID/HCD/ETD triplet fragmentation mode on 2hr gradients. Additionally, data of an antigen-specific fraction with the highest protein yield from week 14 of a single rabbit was collected on the Orbitrap Lumos in HCD/EThcD doublet fragmentation mode as stated above. In the peptide immunization, serum was split and purified by either peptide A conjugated to resin or peptide B conjugated to resin. For each peptide purification, the fraction with highest titer differential between peptides in ELISA was selected. MS/MS data was generated on peptide purified fractions using HCD/EThcD doublet fragmentation mode on the Lumos.

### 2.4 Informatics

Transcript sequencing data was processed with in-house pipeline for trimming adaptors, pair-stitching, and filtering of low quality reads. Processed raw sequences were collapsed and error corrected using IgRepertoireConstructor [31]. Repertoire sequences were V(D)J labeled using alignment [4] to rabbit gene references from IMGT. V(D)J labeling reports complementarity-determining regions (CDRs), and sequence differences from gene references (i.e., somatic hypermutation events). Repertoires construction, V(D)J labeling, CDR identification, and mutation calling was performed independently on each sample.

MS/MS data was collected for each antibody chain and enzyme digestion. Each run was searched against a non-redundant database of transcript sequences with minimum RNA abundance of 2 across all chain-specific repertoires from bleeds and tissues, plasma contaminant proteins, and digestion enzyme sequences using MSGFDB [18]. MSGFDB was run in merge fragmentation spectra mode with high accuracy error thresholds (20ppm precursor m/z error), and ETD model was used to search and score EThcD spectra since no EThcD models have been trained. To estimate false discovery rate (FDR), the target-decoy approach was applied by reversing all sequences from the target database. Peptide spectrum matches surpassing a 1% estimated FDR were reported. Assignment of proteomic clone presence in serum is determined by 100% peptide coverage of CDR3, 75% unique peptide coverage of CDR3, and at least one antibody sequence of the clone contains 100% antibody coverage. Clones without *clone-unique* peptide coverage were not included as observed in serum. Additionally, for candidate monoclonal antibody discovery, heavy and light chain clones were selected based on strength of unique proteomic coverage.

Additionally, MS/MS data not matched to the database was searched for mutations de novo. Briefly, raw spectra were deconvoluted to remove isotopic envelopes and de-charge fragment ions, and prefix residue mass (PRM) spectra were generated by training a predictive model on datasets from [30, 34, 28] and procedures motivated by [9]. PRM spectra were then aligned to a database of theoretical PRM spectra generated from repertoire-matched unique peptides allowing for a single amino acid difference in the alignment. See S.I. Appendix H for details on alignment scoring and benchmarks on mutation differences between peptide spectrum matches. Mutated peptide spectrum matches were re-scored with spectral probability [18], required to have a minimum spectral probability of 10^−8^, and not be found in sequenced repertoires. Antibody mutations were called by mapping mutated peptides to repertoire sequences. Mutations for isoleucine (Ile) and leucine (Leu), asparagine (Asn) to asparatic acid (Asp), and glutamic (Glu) acid to glutamine (Gln) were excluded since parent mass differences are zero or identical to deamidation post-translational modification.

### 2.5 Antibody expression and validation

Candidate antibody sequences were picked from 11 different heavy and 9 different light chain clones for the recombinant Beta-gal immunization. Heavy and light chains were synthesized and cloned into respective rabbit IgG1 and IgK vectors (Absolute Antibody) with Sanger DNA sequence confirmation. All pairwise combinations of heavy and light chain vectors were transfected into HEK293 cells at a 2mL scale and cultured for six days. Crude supernatant from the hundred transfections were protein A purified and run on SDS-PAGE gel to confirm antibody expression. For both the recombinant Beta-gal and Beta-gal peptide immunizations, Protein A purified supernatant was tested to confirm antibody binding with indirect ELISA. Positive control anti-Beta-gal antibody (Ab00135-23.0 from Absolute Antibody) and PBS blanks were also tested. HRP conjugated goat anti-rabbit IgG secondary antibody (5196-2504, Bio-rad) was used for detection. Kinetic analysis of purified antibody was performed on Octet Red96 (Forte Bio) following vendor protocol.

## 3 Results

### 3.1 Ambiguity in determining clone antigen-specificity exclusively from repertoire sequencing

Two rabbits were immunized with recombinant Beta-gal with boosts at weeks 3, 6, and 10. New B-cell clones specific to Beta-gal are expected to emerge upon immunization and pre-existing clones are expected to expand with more mature antibodies after boosts. To track the progression of response to Beta-gal, Ig transcripts were sequenced from PBMC at weeks 0-14 (see Figure 1 for read counts and weeks) and spleen and bone marrow at week 14. Antibody transcripts that share an exact CDR3 are presumed to originate from the same progenitor B-cell clone, and their abundance can be tracked across time points. From each time-point sample, heavy and light chain repertoires were sequenced at consistent depths. For heavy chain, the raw sequencing depth for each sample is on average 61, 227 ± 8, 855 standard deviation and after filtering and error correction is 22, 662 ± 4, 478. Light chains repertoires were sequenced to the same raw depth, 78, 490 ± 8, 146, after filtering and error correction, average depth is 6, 534 ± 1, 229. The repertoires have similar features to previous reports on hyperimmunized rabbits. The average variable heavy chain CDR3 length prior to immunization is 15.2 ± 3.1 and post-immunization is 14.7 ± 2.9, which is greater than the 12.0 − 12.3 range reported by [19], but consistent with 14.8 ± 3.6 reported by [22]. The average variable light chain CDR3 lengths for each immunization is also consistent with pre-immunization 11.6 ± 1.3, and post-immunization 11.7 ± 1.7(see S.I. Appendix S2). Also, somatic diversification of antibodies is similar between repertoires across time points and tissues. The average somatic hypermutation observed for compartments ranged between 10.6 − 11.6 for heavy chain transcripts, and 14.9 − 15.7 for light chain transcripts, which is consistent to averages of 9.8 − 12.4 and 14.9 − 15.7 observed by [19]. The repertoire constructed from bone marrow shows a higher rate of somatic hypermutation compared to other repertoires for both heavy and light transcripts (see S.I. Appendix S3). From bulk B-cell sequencing, discerning which heavy and light chain transcripts generated in response to an antigen is challenging, and typically requires sequencing approaches that pair chains or cell sorting [12]. From the repertoire analysis alone, there is no obvious B-cell clone that was generated and expanded in response to Beta-gal. There are 2, 115 ± 783 heavy chain clones appearing for each post-immune repertoire, which amounts to 3, 262 clones that appear in at least two post-immune repertoires. Figure 1b shows the variable heavy chain clones as a fraction of the repertoire sampled at each time point. No individual clone showed expansion in transcript abundance or unique antibody sequences in progression of the Beta-gal immunization and boosts.

### 3.2 Repertoire clone selection from affinity purified serum IgG

Total serum IgG from the recombinant Beta-gal immunized rabbits contain a complex mixture of antibodies, of which a small fraction may bind to Beta-gal. To better characterize the IgG response to Beta-gal, the final fraction of serum underwent affinity purification against the recombinant antigen using two different methods. In the first approach, the two individuals were pooled and all fractions off the column were also pooled; while the second approach purified each individual separately, fractionated the eluted proteins into six fractions, and the fraction with highest concentration was selected for LC-MS/MS. The pooled sample contains 22,373 triplet (CID/ETD/HCD) spectra, while the fractionated sample contains 59,015 and 51,655 (HCD/EThcD) spectra for heavy chain and light chain, respectively. The increase in spectral count for the fractionated sample is due to the use of a mass spectrometer with a higher scan rate, and not related to sample or instrument method. Spectra are searched against the combined repertoires from weeks 0 to 14 to determine which antibodies were present in the final purified serum.

A limitation of bottom-up MS/MS is distinguishing the presence of individual antibodies. Antibodies circulating in the serum that belong to the same clone (i.e., unique CDR3 sequence) may differ by only a few mutations. In immunoproteogenomic approaches, antibodies are digested into peptides, leading to ambiguity of peptide assignment to a unique antibody sequence. Unique CDR3 sequences have fewer ambiguous peptide assignments and clones can be more easily identified. Wine et al., [39] and Lavinder et al., [22] rely on tryptic digestion of antibody sequences, which result in a peptide uniquely covering the CDR3 to mark the presence of a clone. However, clones could be missed if lysines or arginines are present in the CDR3. Additionally, tryptic peptides covering light chain CDR3s are often too long to be analyzed by MS/MS. Cheung et al. [8] used multiple proteases and required: at least 65% antibody sequence coverage, 12 unique peptides, and 95% CDR3 coverage to determine the presence of an antibody in serum. The use of multiple proteases, as in this study, enables analysis of both heavy and light chains and facilitate selection of individual antibody sequences.

In this study, a clone is defined by a unique CDR3 sequence and is proteomically observed if there is 100% peptide coverage over the CDR3 with at least 75% unique peptide coverage. Since B-cell sequencing shows the presence of clones across time points, the emergence of proteomically observed clones can be tracked (i.e., clone *spawning*). The spawning plots for the heavy chain and light chain of a single rabbit immunized with recombinant Beta-gal are shown in Figure 2a and Figure 2b. The clone spawning for heavy chain reveals that many clones are shared between not only spleen and bone marrow, but also intermediate time points, particularly weeks 4, 7, 9, and 11. A minority of clones, 2.2% heavy and 7.0% light chains, originated from the pre-immune repertoire, and represent antibodies that bound non-specifically during affinity purification. The proteomically observed light chain clones show less CDR3 diversity and belong to 84 clone clusters compared to 26 heavy chain clone clusters (see S.I. Appendix C).

**Figure 2:**
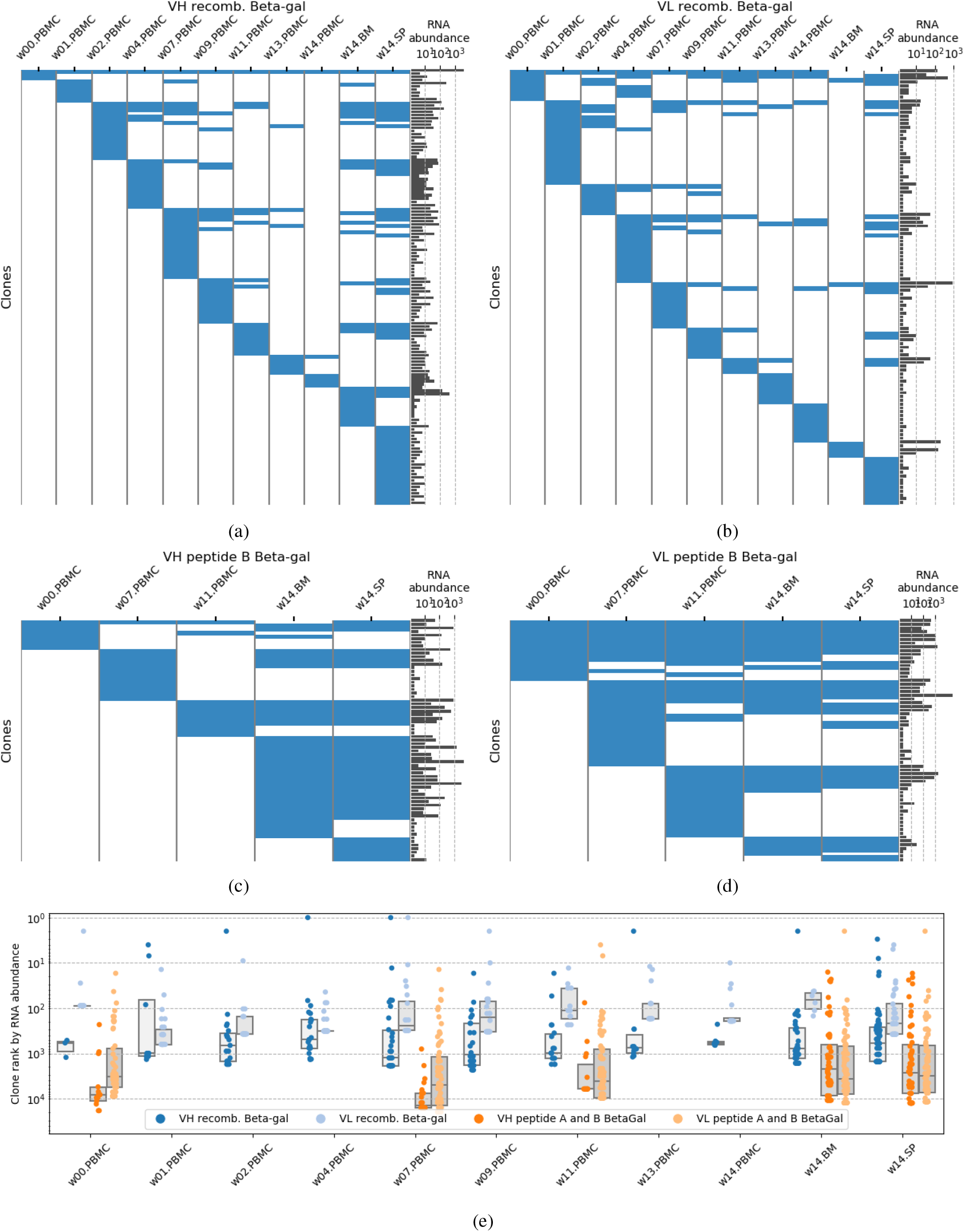
Spawning plots of clones observed Beta-gal purified serum fractions. Clones are identified by unique peptide CDR3 coverage of ≥ 0.75 and complete coverage of the CDR3. There were 136 VH clones in the recombinant Beta-gal purified fraction (a), 115 VL clones identified (b), 67 VH clones in the peptide B purified fraction (c), and 65 VL clones in (d). Log transformed RNA transcript abundance of each clone is shown as a bar plot to the right of spawning plots. e) Scatter plot of proteomically observed clones ranked by RNA abundance within each individual repertoire. Boxes show interquartile ranges of clone rank.

Approximately 27.2% of heavy and 14.8% of light chain clones are *only* observed in bone marrow or spleen and missed in the PBMC; showing just how important those compartments are as a proxy for the serum repertoire. PBMC repertoires had fewer matches to proteomically observed clones, in particular PBMC repertoires in weeks 13 and 14 only contributed 7.4% and 15.7%, for heavy and light chains, respectively. While the B cells were collected at the closest time points to the analyzed affinity purified serum, short-lived plasmablasts generated in response to the final boost at week 10 have limited viability [33] and are unlikely to remain in circulation for 3-4 weeks. The PBMC collections of intermediate weeks 1-11 have a strong match to the affinity purified fraction and account for 65.4% of heavy chains and 69.6% of light chains. Despite spleen and bone marrow being a rich source of proteomically observed antibodies, intermediate PBMC time points provide a reasonable proxy for these compartments to represent the serum.

### 3.2.1 Antigen-specific clones are not the most abundant in repertoires

The abundance of a clone in a repertoire is measured by counting RNA transcripts across all antibodies sharing the clone CDR3. In hyperimmunized animals, highly abundant clones are expected to target the immunogen. As shown in Figure 2e, the clones found in the Beta-gal purified fractions are not the most abundant clones in each Rep-seq repertoire, an observation also reported by Cheung et al. [8]. In the recombinant Beta-gal immunization, proteomically observed heavy chain clones rank between 1-2,322 on RNA abundance among all clones in each repertoire. Even though the clones may not have the highest abundance, PBMC repertoires from weeks 4, 7, and 9 have a statistically significant higher ranking of proteomic clones (one-sided p-value < 0.05 Mann-Whitney U test). Similarly, clones ranked between 2-1,639 and 3-1,510 in bone marrow and spleen, respectively, with only spleen showing a statistically significant enrichment. While spleen and bone marrow are the most comprehensive single sources of proteomically observed clones, the abundance rank in bone marrow and spleen is lower than intermediate PBMC repertoires, when the clone is present in both. On average, RNA abundance rank of proteomically observed light chains are higher than heavy chain, however this is attributable to lower diversity of light chain clones. Only repertoires from weeks 7, 9, 11, and 13 show a statistically significant higher abundance for proteomically observed light chain clones (one-side p-value < 0.05 Mann-Whitney U test). However, similar to heavy chain, light chain clones have higher abundance rank in intermediate PBMC repertoires than bone marrow and spleen. The variable presence and sometimes low abundance of proteomically observed clones across repertoires suggest that the most transcriptionally active clones sampled in Rep-seq do not represent the serum immune response to Beta-gal.

### 3.2.2 Affinity purified serum IgG has limited concordance to total IgG

Serum from pre-immune and weeks 1, 7, 11, and 14 were purified with protein G to obtain a total IgG fraction (IGG) to match to repertoires, and determine if affinity purified serum (AP) is necessary to find Beta-gal binding clones. Since both fractions originate from the same hyperimmunized animals, the IGG fraction should comprise of the same antigen-specific antibodies as AP and few non-specific antibodies. The AP and IGG fractions are compared at various levels: peptide IDs, Ab coverage, and coverage of CDR3 clone sequence. At the peptide level, a high degree of overlap is necessary to identify the same antibodies. However, the two fractions had only 30.1% of unique peptide identifications in common (see Table S5).

The low recovery of peptide IDs in the IGG fraction could be due to the presence of numerous antibodies that are not specific to the antigen. However, the IGG fraction also has higher noise, as evidenced by the 20% higher relative level of peptides mapping only to the host organism proteome (1564, or 63%, for IGG versus 848, or 42%, for AP). Approximately one third (501 of 1564, 20% of IGG total) of host-mapping peptides in IGG mapped to serotransferrin. While IGG was treated with protein G to isolate IgG, this high abundance serum protein was also retained. At least one other study of serum antibodies report serotransferrin as a common contaminant [10].

While a considerable percentage of identified peptides in the IGG fractions are to contaminants, tracking heavy chain clones by CDR3 sequences could still be performed. Table S6 shows the number of unique CDR3 sequences with different threshold requirements on proteomic coverage. While no clone has complete coverage of the CDR3 in the IGG fraction, unlike the AP fraction, we can still track a handful of clones at a reasonably high coverage of 85%.

Proteomic tracking of clones can provide insight into how serum antibodies either persist across the immunization; or if new clones are created after each boost of the antigen. Figure 3 shows clones with peptidic evidence occurring in one or more time points. Each node represents a single clone at a particular time point. Two identical clones are joined by an edge if they co-occur in two different time points. To accommodate for lower depth of proteomic sampling, clones with peptidic evidence covering at least 90% of the CDR3 region are considered as existing in the serum for each time point.

**Figure 3:**
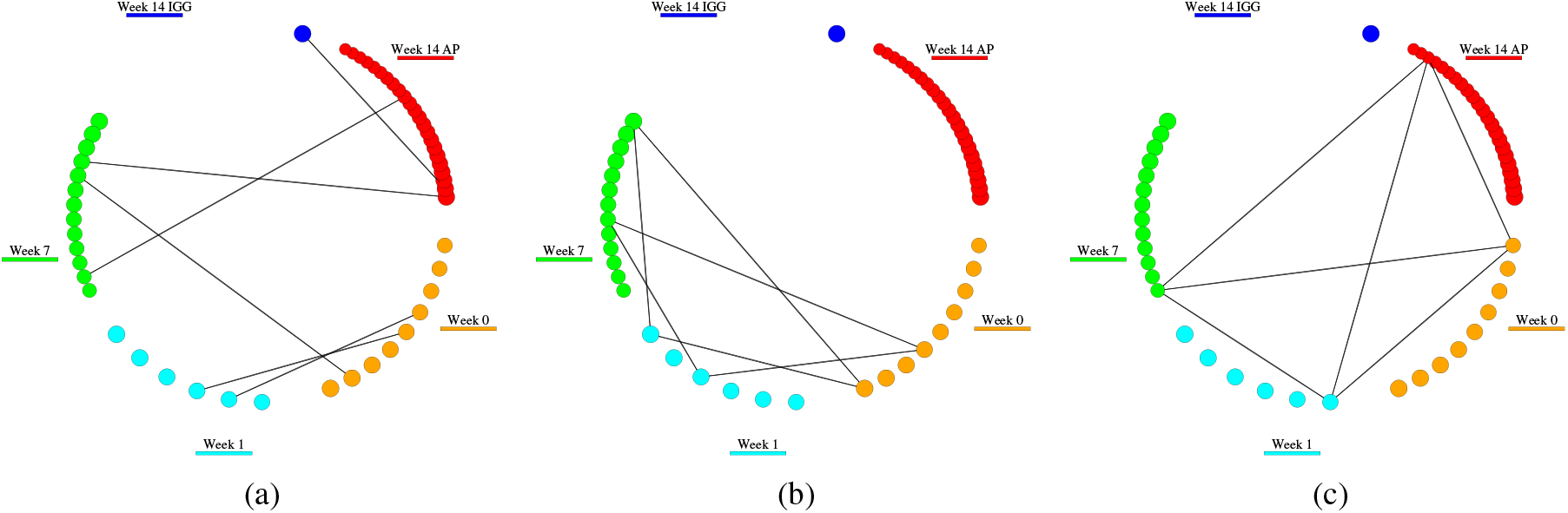
Proteomically represented clones. Clones at fractions from different time points are shown around each circle, and an edge is drawn between two nodes if that clone was present in both time points. Clones with at least 90% of their CDR3 is supported by one or more peptide are shown. Clones co-occurring in two (a),three (b),and four (c)fractions are shown separately.

Figure 3 shows that most clones at week 1 are shared at week 0, suggesting that the antibodies raised against Beta-gal have not yet become prominent in the serum. At week 7, serum has high antibody titers to Beta-gal, as measured by ELISA (data not shown). The week 7 IGG fraction has 13 identified clones where 3 clones are shared with the terminal AP fraction and 4 are shared with week 0 or week 1, showing week 7 production bleeds still contain a combination of pre-immune antibodies and antibodies to Beta-gal.

The majority of the terminal AP fraction clones, 20 of 23 clones, are specific to that fraction alone. While the terminal AP fraction was enriched for antibodies against Beta-gal, very few are shared with other non-terminal weeks, suggesting that these clones did not exist prior to the immunizations. Relaxing the peptide evidence required for determining the existence of a serum clone to 85% shows that 4 of 5 clones in the IGG terminal fraction are also present in the AP fraction (see Figure S6). This evidence suggests that the IGG fraction is a similar serum repertoire to the AP fraction, but with fewer observed clones.

The comparison of clones identified in AP and IGG fractions underscores the complexity of proteomic analysis of total IGG. Less stringent purification is prone to more contamination and as a result, lower clone identification rates. The final week 14 IGG fraction had the highest level of host protein and serotransferrin contamination (see Table S5) and lowest number of identified clones. However, affinity purification is also insufficient to entirely exclude contamination. While most of the clones in the AP fraction appear in post-immune repertoires (see Figure 2a), one clone in Figure 3c is present in IGG fractions across weeks and the AP fraction, and is likely in high abundance in serum and non-specific to Beta-gal.

### 3.2.3 Recombinant protein immunogen illicits more diverse immune response than peptide immunogens

Recombinant antigens consist of many epitopes, which could correlate with more diverse B-cell responses and more complex repertoires of circulating antibodies in serum. The immunoproteogenomic approach was repeated to profile the circulating serum repertoire of a rabbit immunized with two peptides Beta-gal portions conjugated to KLH. Rabbits were subjected to the same immunization schedule, and repertoires were sampled at weeks 0, 7, 11 from PBMC and weeks 14 from bone marrow and spleen (see Figure 1). Each repertoire was sequenced more deeply than the recombinant Beta-gal immunization, with 1, 392, 029 ± 138, 094 reads per heavy and light chain repertoire. The heavy and light CDR3 length distribution for the peptide immunization is similar to the repertoires from the recombinant immunization, post-immunization mean length being 14.0 ± 2.6 and 11.9 ± 1.7, respectively (see S.I. Appendix S2). Additionally, somatic antibody diversification is similar with 10.3 − 12.0, and 14.8 − 15.9 mean mutations per repertoire for heavy and light chain respectively. There are 29, 580 ± 2, 371 heavy chain clones appearing for each peptide immunization repertoire, which amounts to a total of 34, 268 clones that appear in at least two post-immune repertoires. Similar to the recombinant Beta-gal immunization, there are no obvious features to easily determine which antibodies from the repertoires bind to Beta-gal.

To find Beta-gal binding antibodies, the final bleed at week 14 was purified into separated fractions by affinity to peptide A and B. The rabbit generated a stronger response to peptide B than peptide A, as measured by higher antibody titers to peptide B in ELISA at both week 7 and 14 serum collections (data not shown). Serum from week 14 was affinity purified against each peptide separately, with the highest titer fraction being reduced into heavy and light chains and subjected to multi-enzyme digestion and LC-MS/MS. A similar number of spectra was generated for each chain as the final recombinant Beta-gal affinity purification (see Table S1). After searching the spectra against the peptide immunized rabbit repertoires, only 20 heavy and 4 light chain clones are proteomically present in the peptide A purified fraction, and 67 heavy and 65 light chain clones present in the peptide B fraction.

The proteomically observed clones in the peptide fractions did not have the highest RNA abundance among repertoires, similar to the clones identified in the recombinant Beta-gal affinity purification. The heavy chain clones observed across both peptide immunization fractions appeared in RNA abundance ranks 16 to 12,451 in bone marrow and spleen, as shown in Figure 2e. Unlike the recombinant Beta-gal immunization, heavy chain clones in the peptide purified fractions rank higher in bone marrow and spleen than PBMC repertoires. Only spleen and bone marrow have a statistically significant higher RNA abundance ranks of proteomically observed heavy chain clones over all Rep-seq clones (one-side p-value < 0.05 Mann-Whitney U test). Light chain clones had a similar distribution of RNA abundance ranks between repertoires, with proteomically observed clones significantly enriched for in each tissue and time-point. Even though proteomically observed clones have higher RNA abundances than other clones, the majority of proteomically observed clones still have abundance ranks below 100. The affinity purification narrows to observing clones to one specific peptide, however there are other epitopes on the carrier protein and background that contribute to the more abundant antibodies in the B-cell repertoires.

Similar to the recombinant immunization and purification, the spawning of proteomically observed clones in the peptide B purified fractions shows most clones are observed in spleen and bone marrow (see Figure 2c and 2d). Due to the deeper sequencing depth, more clones are present in multiple samples, e.g., of the 54 heavy chain clones observed in spleen 83.3% are also in bone marrow. Although only two post-immune PBMC repertoires were sampled, the PBMC repertoires contribute to 38.3% of heavy and 84.0% of light chain clones observed proteomically in serum. The week 7 PBMC repertoire is particularly important and uniquely contributes to 15.1% of heavy chain clones compared to 4.6% for week 11 and 9.3% for spleen.

The fewer epitopes on peptides resulted in 36.0% fewer heavy chains clones in peptide A or B purified fractions compared to the recombinant Beta-gal purified fraction. The reduced diversity is accentuated to ≈70% fewer clones when restricting identified heavy chain clones to require at least 90% and 99% unique peptide coverage of CDR3s as shown in Figure S5. The reduction in light chain clone diversity between immunizations is similar, ranging from 27.1% to 40.0% depending on the criteria for clone identification (see Figure S5). The number of heavy and light chain clones is consistent for each fraction, with 65-67 clones found in the peptide B purified fraction, and 115-136 clones found in the recombinant Beta-gal purified fraction.

### 3.2.4 Confirmation of Beta-gal affinity for individual monoclonal antibodies

Antigen-specific candidates were selected for expression and validation based on strong proteomic evidence from the affinity purified samples, and requiring clones to be observed in spleen. Nine different heavy chain clones and ten light chain purified against recombinant Beta-gal were selected. One full length antibody sequence representative was selected for each clone, with the exception of one heavy chain clone where two full-length antibodies with strong peptide support were selected. All pairwise combinations of 10 heavy and 10 light antibody sequences were cloned into a rabbit IgG1 vector and expressed in HEK293 cells. Of the 100 initial candidates, three candidates show binding to recombinant Beta-gal by ELISA (see Figure S7) and were selected for scaled up production and kinetics analysis. Table 1 shows Kd values of these candidates compared to a positive control mAb from [27]. The two picomolar binding antibodies with better affinity than the positive control, share the same light chain sequence. One heavy chain clone appears at RNA abundance ranks 659 and 8 in bone marrow and spleen, and the other heavy chain clone appears at rank 235 in PBMC collected at week 11. The light chain clone appears in PBMC week 4 and 7, and spleen with abundance ranks ranging from 11 to 175. The variable presence and ranking across repertoires of the validated monoclonal antibodies suggests sampling of multiple time-points is necessary to ensure capture of high affinity candidates.

**Table 1:** Kinetic analysis of recombinant Beta-gal candidates compared to positive control from [27].

| Ab | KD (M) |
| --- | --- |
| Ab 1 | 9.7E-11 |
| Ab 2 | 9.6E-11 |
| Ab 3 | - |
| Control | 1.2E-09 |

Similarly, 11 heavy and 9 light chain clones purified against peptide B Beta-gal were selected for expression and binding analysis. The resulting 99 antibodies were expressed, and single-spot ELISA was used to test binding activity of supernatant to recombinant Beta-gal. Two antibodies show binding activity within 1000-fold of the positive control antibody. The two antibodies with activity have distinct light chains, but share the same heavy chain sequence. The heavy chain clone appears in PBMC collected at week 11, bone marrow, and spleen with RNA abundance ranks ranging from 76 to 123 in each repertoire. The light chain clones have abundance ranks ranging from 56 to 7808 in repertoires collected at PBMC week 7 and 11, bone marrow, and spleen. Production of more proteomically observed candidate heavy and light chain pairs from both immunizations would contribute to finding correct pairings, and lead to validation of more high-affinity binders to Beta-gal.

### 3.2.5 The hidden repertoire: antigen-specific antibodies that lie beyond the repertoire

While B-cell sequencing allows for more sensitive interpretation of mass spectra, antibodies in serum without a corresponding B-cell transcript will be missed. To determine this hidden repertoire of unidentified spectra, a spectral networking approach [1] is applied to identify spectra corresponding to antibody mutations. Briefly, spectra are aligned to repertoire mapped peptides allowing for a single amino acid substitution. The alignment reports the optimal amino acid substitution and position on the peptide originating from an antibody in the repertoire. To maintain a consistent scoring scheme as repertoire database search, the un-identified spectra are re-scored with the discovered mutated peptides using spectral probability as calculated by MS-GF [18]. Mutated peptide spectrum matches are then filtered by minimum spectral probability and to ensure no exact match to any portion of Rep-seq sequences.

In the recombinant Beta-gal purified fraction, there are 529 mutated peptides identifications, which contributes to 5.8% and 9.9% more heavy and light chain peptide identification than found in Rep-seq alone. The peptide A Beta-gal and peptide B Beta-gal purified fractions have 75 and 153 mutated peptide identifications, which represents a 5% increase in peptide identifications over Rep-seq alone.

The discovered mutations are assigned to repertoire antibodies and categorized by CDR and framework regions on variable heavy and light domains. Figure 4 shows example mutations to Rep-seq clones along with support from mutated CDR3 peptide spectrum matches. In some cases, a mutated peptide is shared between clones and antibodies, and the corresponding mutation cannot be unambiguously assigned to a single antibody region. Counting mutations using shared mutated peptides is confounding as the mutation could affect one antibody or potentially hundreds. However, 247 mutations could be uniquely assigned to antibody regions, and the discovered mutation counts across different affinity purified fractions: recombinant Beta-gal, peptide A Beta-gal, and peptide B Beta-gal are shown in Table 2. The variable light domains in the recombinant Beta-gal purified fraction have the most mutations compared to variable heavy domain, or peptide purified fractions. Even though the light chain repertoires have the same sequencing depth as heavy chain repertoires, the clone diversity is lower in Rep-seq and likely requires deeper sequencing to capture the same number of clones as heavy chain. The peptide immunization and purifications show similar numbers of variable heavy and light domain mutations suggesting the sequencing depth is sufficient for both.

**Table 2:** Mutation counts not present in Rep-seq and discovered in week 14 affinity purified fractions. Shared mutated peptides can be shared between antibody or clone sequences. Unique peptides are exclusively assigned to antibody sequences.

| Beta-gal purification |  | variable heavy domain |  |  |  | variable light domain |  |  |  |
| --- | --- | --- | --- | --- | --- | --- | --- | --- | --- |
|  |  | cdr1 | cdr2 | cdr3 | fw | cdr1 | cdr2 | cdr3 | fw |
| recomb. | shared mutated peptides | 5 | 14 | 2 | 64 | 29 | 27 | 20 | 180 |
|  | unique mutated peptides | 7 | 5 | 13 | 31 | 15 | 31 | 33 | 53 |
|  | regions with unique mutated peptides | 4 | 3 | 11 | 20 | 12 | 19 | 23 | 37 |
|  | unique mutations | 5 | 4 | 13 | 27 | 14 | 28 | 30 | 52 |
| peptide A | shared mutated peptides | 1 | 2 | 0 | 18 | 1 | 4 | 3 | 18 |
|  | unique mutated peptides | 0 | 0 | 0 | 9 | 3 | 2 | 1 | 13 |
|  | regions with unique mutated peptides | 0 | 0 | 0 | 5 | 3 | 2 | 1 | 11 |
|  | unique mutations | 0 | 0 | 0 | 8 | 3 | 2 | 1 | 13 |
| peptide B | shared mutated peptides | 1 | 12 | 7 | 35 | 7 | 3 | 4 | 36 |
|  | unique mutated peptides | 1 | 6 | 4 | 10 | 1 | 4 | 4 | 18 |
|  | regions with unique mutated peptides | 1 | 5 | 3 | 9 | 1 | 4 | 4 | 13 |
|  | unique mutations | 1 | 6 | 4 | 10 | 1 | 4 | 4 | 17 |

**Figure 4:**
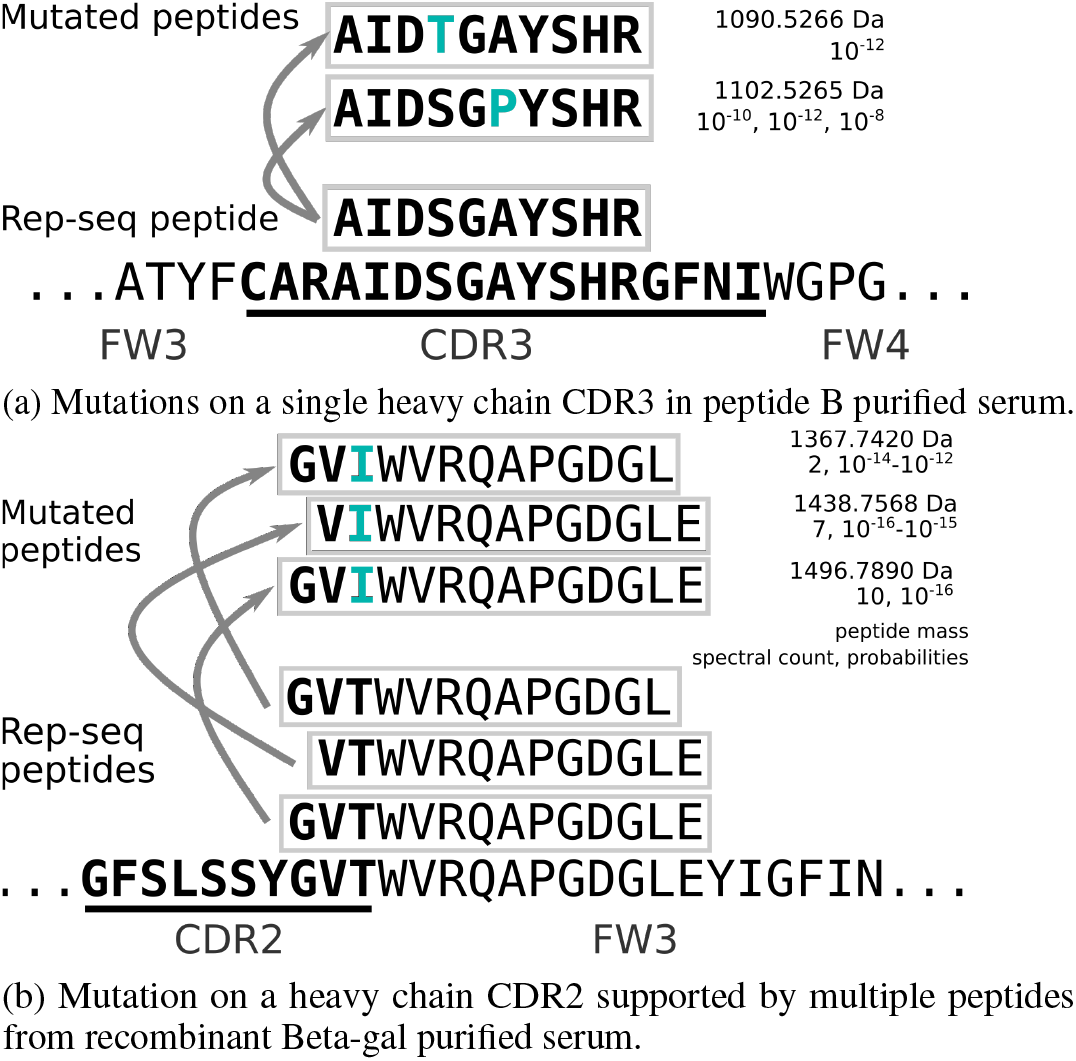
Mutated peptides mapping uniquely to variable heavy domain complementarity-determining region (CDR). Mutations are shown in blue, amino acids overlapping CDRs are in bold, and framework region (FW) is in plain text. The peptide mass, spectral count, and spectral probabilities from supporting peptide spectrum matches are shown next to each mutated peptide.

Unique mutations in the CDR regions are particularly important, as they are the most likely to directly affect binding affinity. There are 54 unique mutations observed across all CDR3s not present in Rep-seq. Considering the CDR3 sequence is used to define clones, there are at least 9.6% and 10.4% more heavy chain clones than found by matching to Rep-seq repertoires from the recombinant Beta-gal and peptide B Beta-gal affinity purified fractions, respectively. As the peptide immunized rabbit did not produce a strong response to peptide A, no new CDR3 mutations are in the peptide A purified fraction. Since the mutation search approach is limited to only single amino acid substitutions from observed repertoire peptides, the number of missed antibodies is a lower bound to unique antibodies present in the circulating serum.

## 4 Discussion

Rabbit polyclonal and monoclonal antibodies are routinely generated for a variety of immunoassays, due to their higher affinity and specificity to unique antigens compared to those derived from other species. Depending on the assay requirements and the availability or cost of the target antigen, antibodies are generated by immunizing rabbits with linear peptides derived from the protein antigen, or either the recombinant or native form of the protein. We applied an immunoproteogenomic approach to profile the polylconal antibodies circulating in serum against two types of targets; recombinant Beta-gal, and two peptides from Beta-gal conjugated to KLH. By profiling the serum proteomics, recombinant Beta-gal results in 52.9% higher diversity of heavy chain clones, compared to the two peptides. The result is not surprising, as there are likely many more epitopes available to generate an immune response on the 116.3kDa recombinant Beta-gal compared to the 2.2kDa and 2.4kDa peptides. However, investigation in more animals and other species are needed to show the diversity of the polyclonal response correlates with immunogenic epitopes on an antigen.

The immunoproteogenomic approach provides new insight into antibodies circulating in the serum over Rep-seq. In many applications, Rep-seq is used to profile the immune response to a specific molecule. While Rep-seq provides a comprehensive view of full-length heavy and light chain sequences, after analyzing B-cell repertoires across time points and different tissues, there is no obvious feature that can be used to discriminate specificity to Beta-gal from Rep-seq alone. Serum purification for Ig with affinity to Beta-gal and MS/MS analysis highlights Ig sequences likely generated by the immunized rabbit to respond to Beta-gal. Despite sequencing depths of 70,000 to 1,390,000 reads per repertoire, antibodies circulating in serum and specific to a target antigen of interest do not always appear in each repertoire. During the immunizations, Rep-seq performed on PBMCs collected throughout the immunization schedule to show when antibody clones originate. Spleen is expected to be the richest source of antibodies [26], and indeed Rep-seq repertoires from spleen had the most comprehensive matching of proteomically observed clones, with 41.2%-67.2% heavy and 26.1%-50.7% light chain of all proteomic clones appearing in spleen. Additionally, many of the clones found in spleen were observed in earlier PBMC repertoires, and earlier PBMC repertoires contributed to 47.8%-72.8% of heavy chain clones. The finding suggests high affinity antibodies may arise earlier than typical polyclonal production bleeds at 7 and 14 weeks in rabbits. Efforts to discover polyclonal and monoclonal antibodies in rabbits do not need to sacrifice animals to harvest spleen and bone marrow to find B cells with high-affinity antibodies, since the compartments could be replaced by multiple PBMC samplings.

Analysis of polyclonal antibodies using the immunoproteogenomic approach is limited by capabilities of MS/MS. Firstly, in bottom-up proteomics, the polyclonal antibodies are digested into smaller peptides for MS/MS to generate identifiable peptide spectra. This leads to ambiguous peptide assignments to multiple antibodies in a repertoire, and presence or absence of individual antibodies is difficult to determine. Even determining the presence or absence of clones is challenging, with multiple approaches for clone identification. Wine et al. [39] and Lavinder et al. [23] limited their analysis to tryptic peptides, which simplifies clone identification to only observing peptides that span the majority of heavy chain CDR3s. In this study, multiple digestion enzymes are used to generate more diverse peptides to fully cover antibodies similar to [8], but using a more stringent definition of clone identification. Relying on tryptic peptides alone would have resulted in observing only 61% of recombinant Beta-gal purified clones, and 49% of Beta-gal purified clones, due to presence of R and K cleavage sites within the CDR3s. Secondly, results may change depending on use of affinity or protein A purification to profile the polyclonal antibodies. Protein A purification yields an IgG fraction that is expected to be most analogous to Rep-seq. However, fewer peptides are identified than the affinity purified fractions, with only a single clone appearing in both AP and IGG fractions. The fewer peptide identifications is likely due to serrotransferrin contamination and undersampling of antibodies, which could be circumvented in the future with more extensive purification and sensitive MS/MS optimization. Lastly, while there are a number of approaches for pairing heavy and light chains in Rep-seq experiments, albeit at a lower throughput than bulk B-cell Rep-seq, native chain pairing is lost in bottom-up proteomics. To validate specificity of affinity purified and proteomically observed clones, we selected a subset of candidate heavy and light chains from the recombinant and peptide Beta-gal experiments to co-express as monoclonal antibodies. Three and two of the expressed antibodies from the recombinant Beta-gal and peptide B Beta-gal candidate selection bind to recombinant Beta-gal as confirmed by ELISA. Of the three binders from recombinant Beta-gal, further validation showed two had higher affinity (10^−11^*M* Kd) than the control Beta-gal antibody. Lack of binding from other candidates is likely due to incorrect heavy and light chain pairing.

Rep-seq has largely been used as a proxy for gauging the immune response, however our findings, like others [8, 32] show highly specific antibodies observed in serum are not the most transcriptionally abundant in repertoires. For example, in spleen, most proteomically observed clones for either chain rank between 100 to 10,000 in RNA abundance. Additionally, proteomically observed antibodies are still missed in intermediate Rep-seq collections. The matching of Rep-seq to antibody proteins depends on many factors including: timing of collecting samples, B cell sampling, location/compartment of sampling, and bias of primer set. We also report findings of additional peptide sequences that did not match repertoires using mutation-tolerant search. Based on these findings, the affinity purified serum fractions contained at least 10% more clones than detectable by immunoproteogenomics approach and potentially many more missed antibodies. The number of missed antibodies is a lower bound on missed clones, as the algorithm is limited to finding missed antibodies related by one amino acid change to a Rep-seq transcript and peptide evidence of the transcript. Missing antibodies with entirely distinct V(D)J recombination events would not be found by mutation-tolerant search. Further improvements to *de novo* sequencing of antibodies, would reveal a more complete characterization of antibodies circulating in serum.

## A Datasets

**Table S1:** Number of immunoglobulin sequences collected. Each row shows a different sample, and samples span multiple time points and tissues. VH=heavy chain; VK=kappa light chain; PBMC=perhipheral blood mononuclear cells; BM=bone marrow.

| Week and Tissue | recomb. Beta-gal Rabbit1 |  | recomb. Beta-gal Rabbit2 |  | peptide Beta-gal Rabbit1 |  |
| --- | --- | --- | --- | --- | --- | --- |
|  | VH | VK | VH | VK | VH | VK |
| w00.PBMC | 86426 | 95910 | 78854 | 83501 | 1216587 | 1442682 |
| w01.PBMC | 86686 | 95536 | 82597 | 83992 | - | - |
| w02.PBMC | 84488 | 94238 | 86253 | 80734 | - | - |
| w04.PBMC | 80598 | 91282 | 83718 | 84343 | - | - |
| w07.PBMC | 88982 | 96098 | 88403 | 81720 | 864333 | 1226211 |
| w09.PBMC | 84043 | 93300 | 86056 | 76859 | - | - |
| w11.PBMC | 85285 | 93745 | 86092 | 84765 | 1066980 | 1241201 |
| w13.PBMC | 78991 | 93187 | 76639 | 79735 | - | - |
| w14.PBMC | 77974 | 91031 | 79423 | 87200 | - | - |
| w14.BM | 86381 | 93056 | 64072 | 89826 | 1139749 | 1331333 |
| w14.SP | 82029 | 87250 | 85532 | 87912 | 1370178 | 1259542 |

**Table S1:**
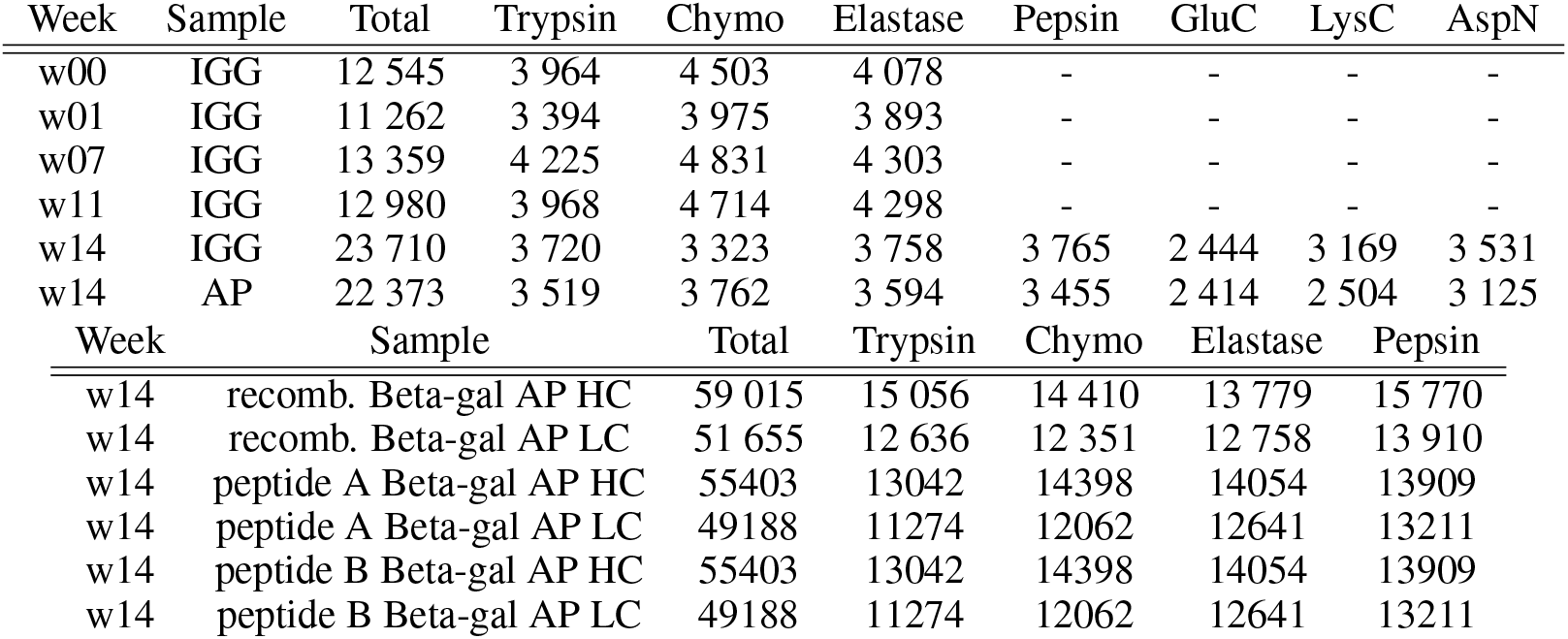
Number of precursors spectra was generated for. Each row shows a different sample, and each column shows the number of precusor ions captured for that enzymatic digest. CID spectra was acquired for each precursor in week 0, week 1, week 7, and week 11 samples. CID, HCD, and ETD fragmentation spectra was generated for each precursor in week 14 AP/IGG samples, and HCD and EThcD fragmentation spectra was generated for each precursor in week 14 HC/LC samples. AP=affinity purified; IGG=IgG purified.

## B Repertoire analysis

**Figure S2:**
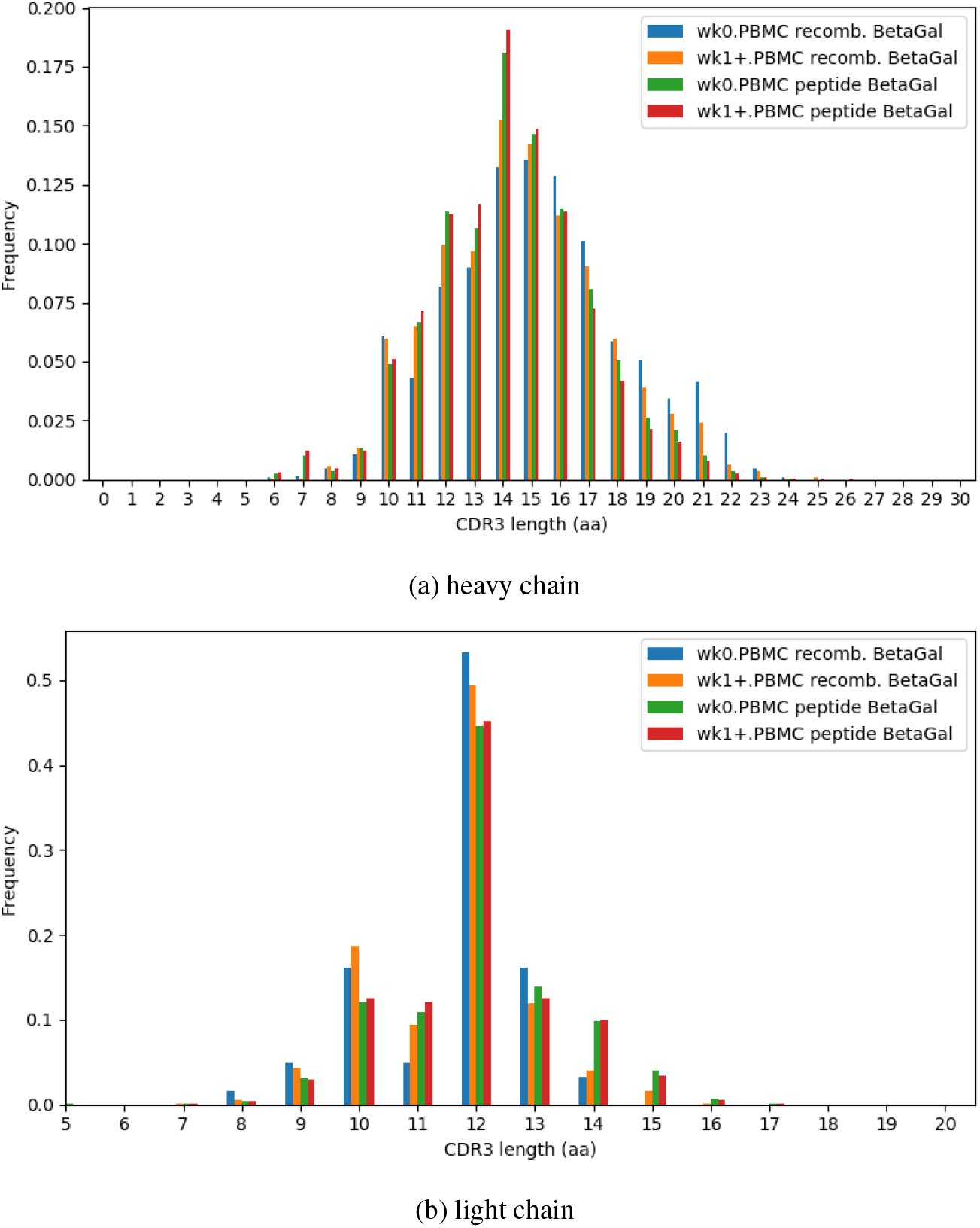
CDR3 length distributions of repertoires separated by pre- and post- immunization. Recombinant and peptide immunized rabbits had similar distributions of CDR3 length for both vh and vk.

**Table S3:** Mean counts of mutations per tissue and time point.

| Week | Tissue | rBeta-gal Rabbit2 |  | pBeta-gal Rabbit1 |  |
| --- | --- | --- | --- | --- | --- |
|  |  | VH | VK | VH | VK |
| 0 | PBMC | 10.6±6.4 | 15.7±4.2 | - | - |
| 1-14 | PBMC | 11.4±5.8 | 14.9±3.7 | 11.4±5.4 | 15.7±4.0 |
| 14 | BM | 12.4±5.8 | 16.4±4.4 | 11.6±5.3 | 16.3±4.5 |
| 14 | Spleen | 11.6±5.2 | 15.1±3.7 | 12.0±5.5 | 16.3±4.5 |

**Figure S3:**
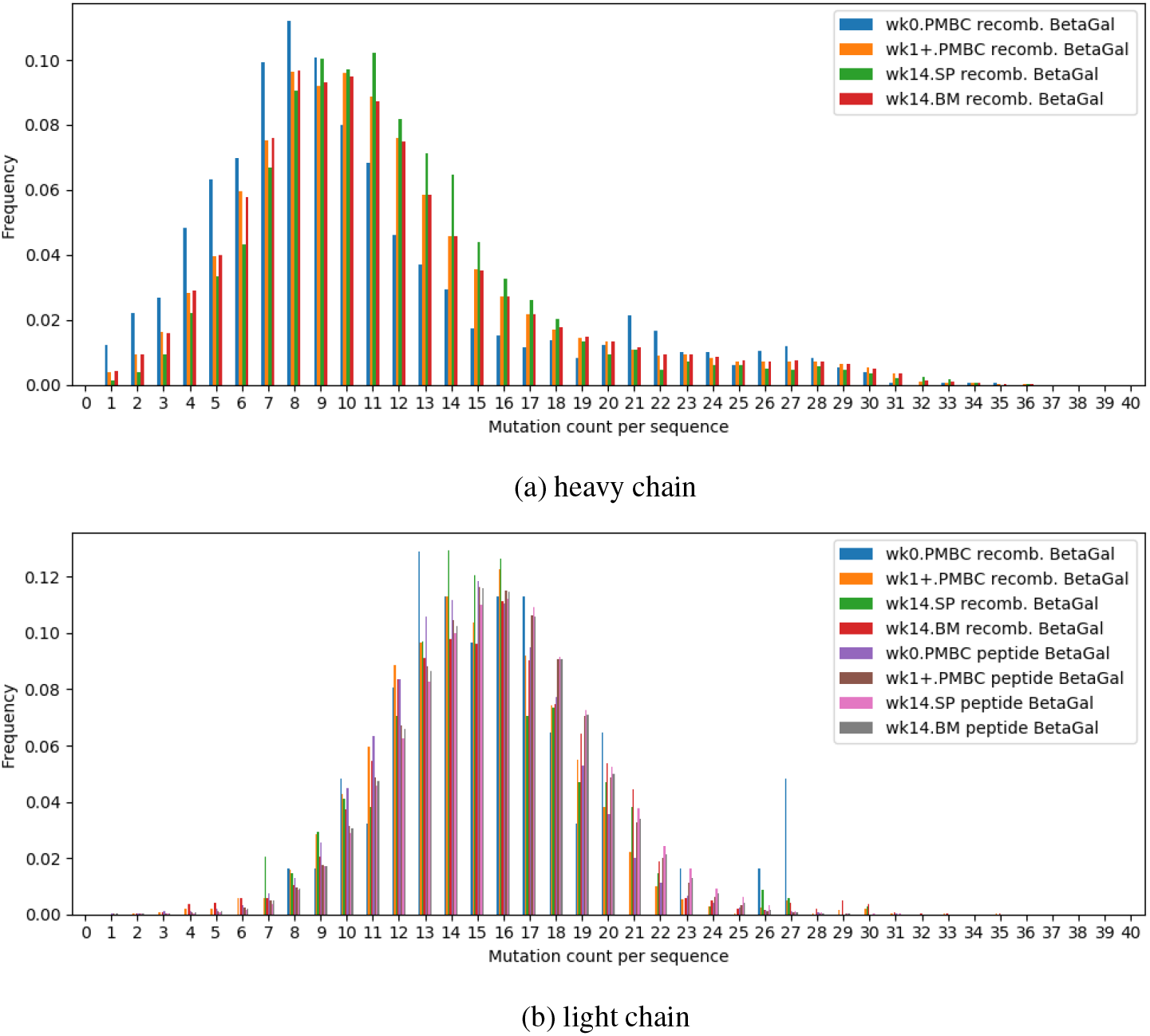
Somatic hypermutation distributions for pre-immune (week 0), post-immune PBMC (weeks 1-14), post-immune SP (spleen at week 14), and post-immune BM (bone marrow at week 14).

## C Proteomically observing clones

**Figure S4:**
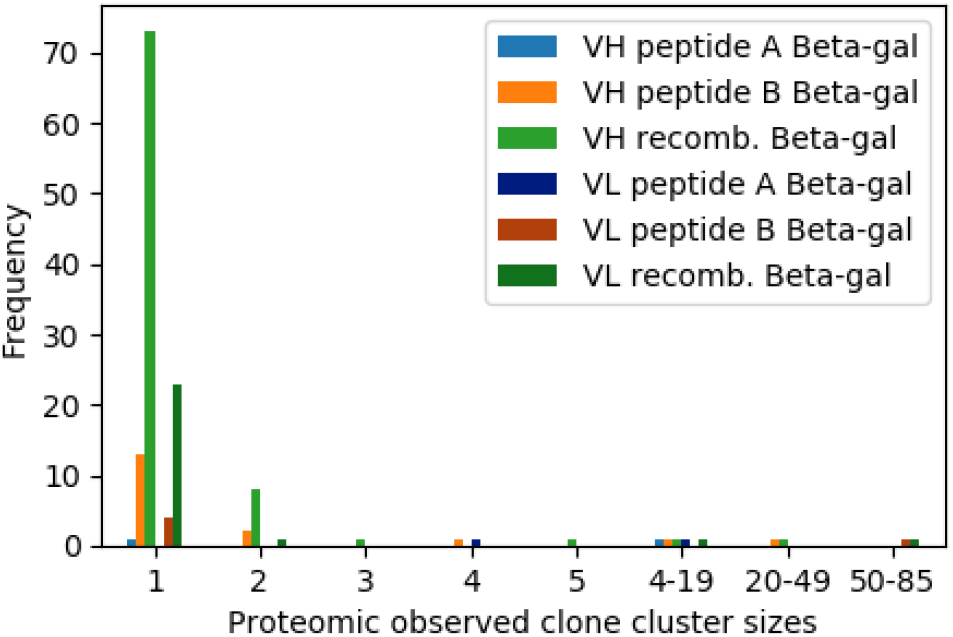
Frequency of proteomically observed clone cluster sizes. Clones are counted as unique cdr3 sequences. Clone clusters were generated from antibody nucleotide sequences using IgRepertoireConstructor [31] at 90% similarity of normalized Hamming distance.

**Figure S5:**
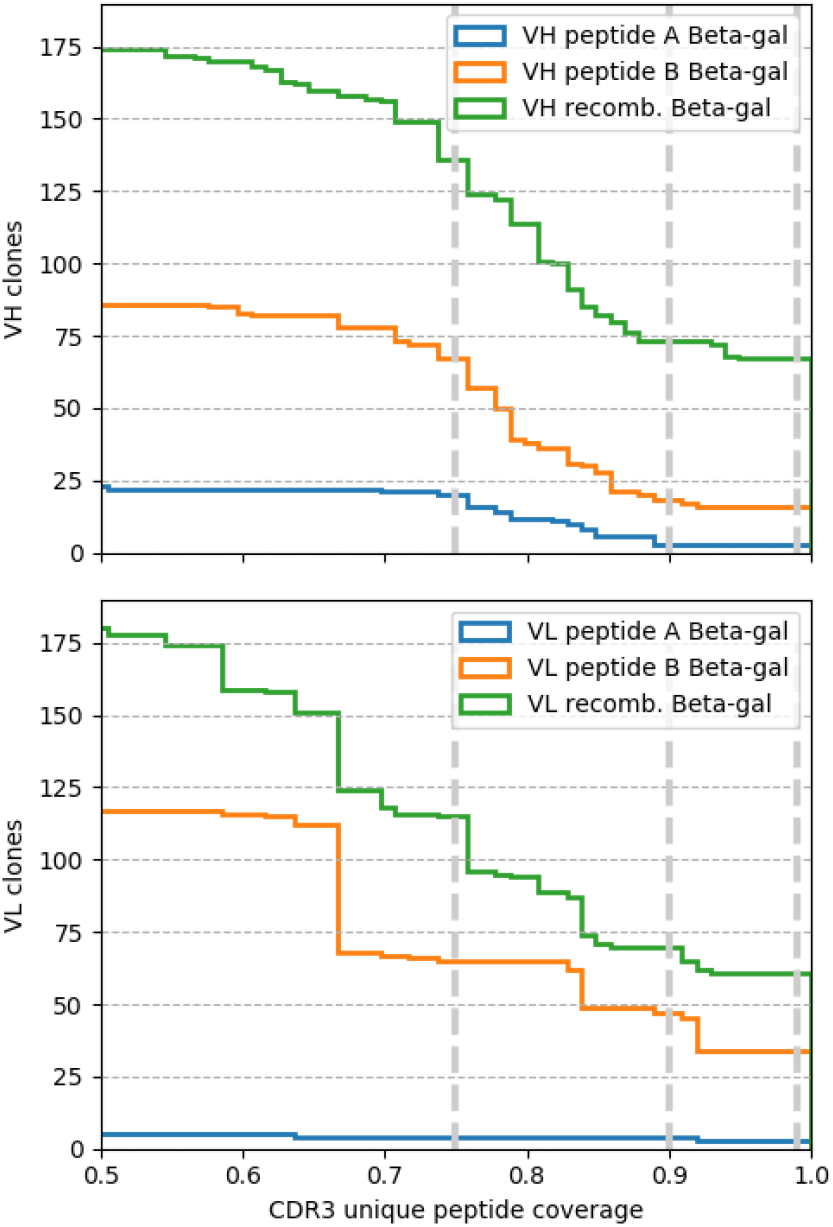
Number of heavy (VH) and light chain (VL) clones at differing thresholds of CDR3 unique peptide coverage. Vertical lines mark coverage thresholds of 75%, 90%, and 99%.

## D Peptide identifications in IGG versus AP in recombinant Beta-gal immunizations

**Table S5:** The number of peptide identifications by sample and enzymatic digest from pooled week 14 serum.

| Set | Total | Trypsin | Chymo | Elastase | Pepsin | LysC | AspN | GluC |
| --- | --- | --- | --- | --- | --- | --- | --- | --- |
| AP | 2007 | 356 | 364 | 313 | 641 | 268 | 277 | 73 |
| IGG | 2474 | 440 | 319 | 351 | 847 | 293 | 334 | 170 |
| $AP \cap IGG$ | 1039 | 199 | 139 | 173 | 333 | 108 | 139 | 47 |

## E Serum contaminants

**Table S5:** Host mapping peptides in the final two samples.

| Sample | # peptides | # host mapping (%) | # mapping to serotransferrin (%) |
| --- | --- | --- | --- |
| AP | 2007 | 848 (42%) | 0 (0 %) |
| IGG | 2474 | 1564 (63%) | 501 (20%) |

While the AP sample has fewer distinct peptides, it has far fewer mapping to the host organism. IGG shows that the dominating host protein contaminant is serotransferrin. At least one other study of serum antibodies report serotransferrin as a common contaminant [10].

## F Clone tracking

**Figure S6:**
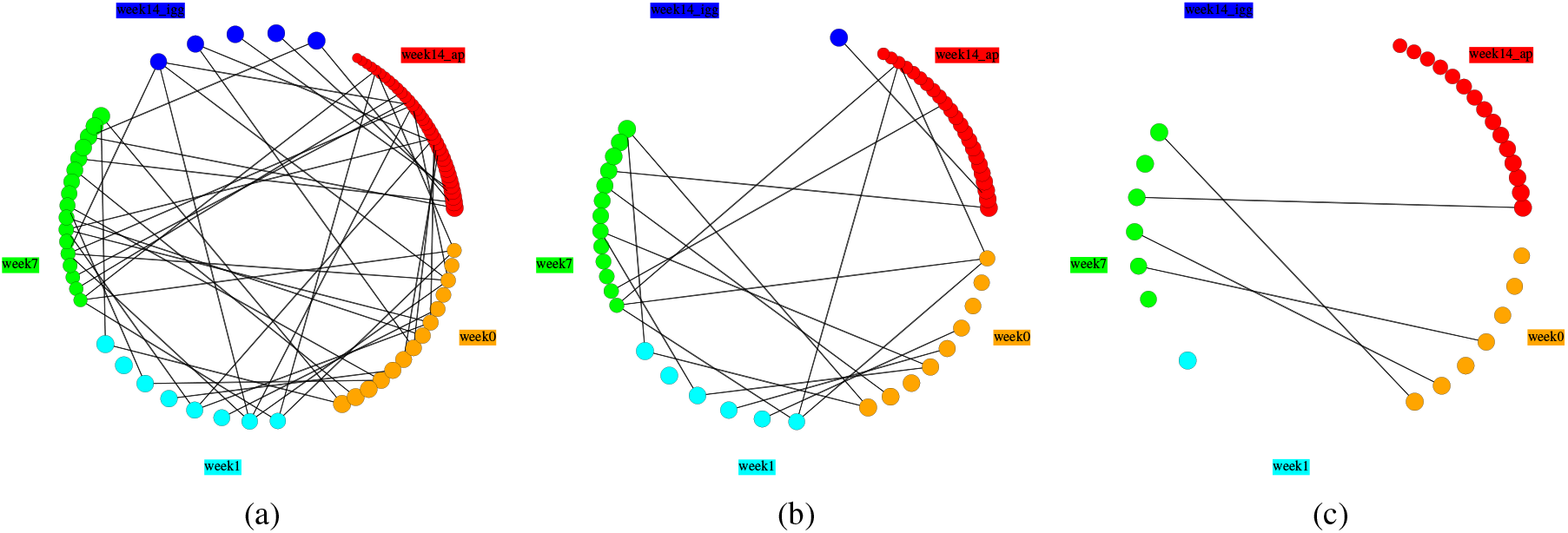
Proteomically represented clones. Clones at different time points are shown around each circle, and an edge is drawn between two nodes if that clone was present in both time points. Clones at 85%, 90%, and 99% peptide coverage are shown in a), b), and c), respectively.

**Table S6:** Clone overlap between AP and IGG samples. The columns show the number of unique CDR3 sequences at different levels of proteomic evidence.

| CDR3 threshold | AP | IGG | $\vert \text{AP} \cap \text{IGG}\vert $ | $\frac{\vert \text{AP} \cap \text{IGG}\vert }{\vert \text{IGG}\vert }$ |
| --- | --- | --- | --- | --- |
| 0.60 | 466 | 285 | 271 | 0.9509 |
| 0.65 | 181 | 83 | 73 | 0.8795 |
| 0.70 | 115 | 45 | 37 | 0.8222 |
| 0.75 | 52 | 21 | 16 | 0.7619 |
| 0.80 | 33 | 9 | 6 | 0.6667 |
| 0.85 | 20 | 5 | 4 | 0.8000 |
| 0.90 | 14 | 0 | 0 | NaN |
| 0.95 | 11 | 0 | 0 | NaN |
| 1.00 | 11 | 0 | 0 | NaN |

## G Validation

**Figure S7:**
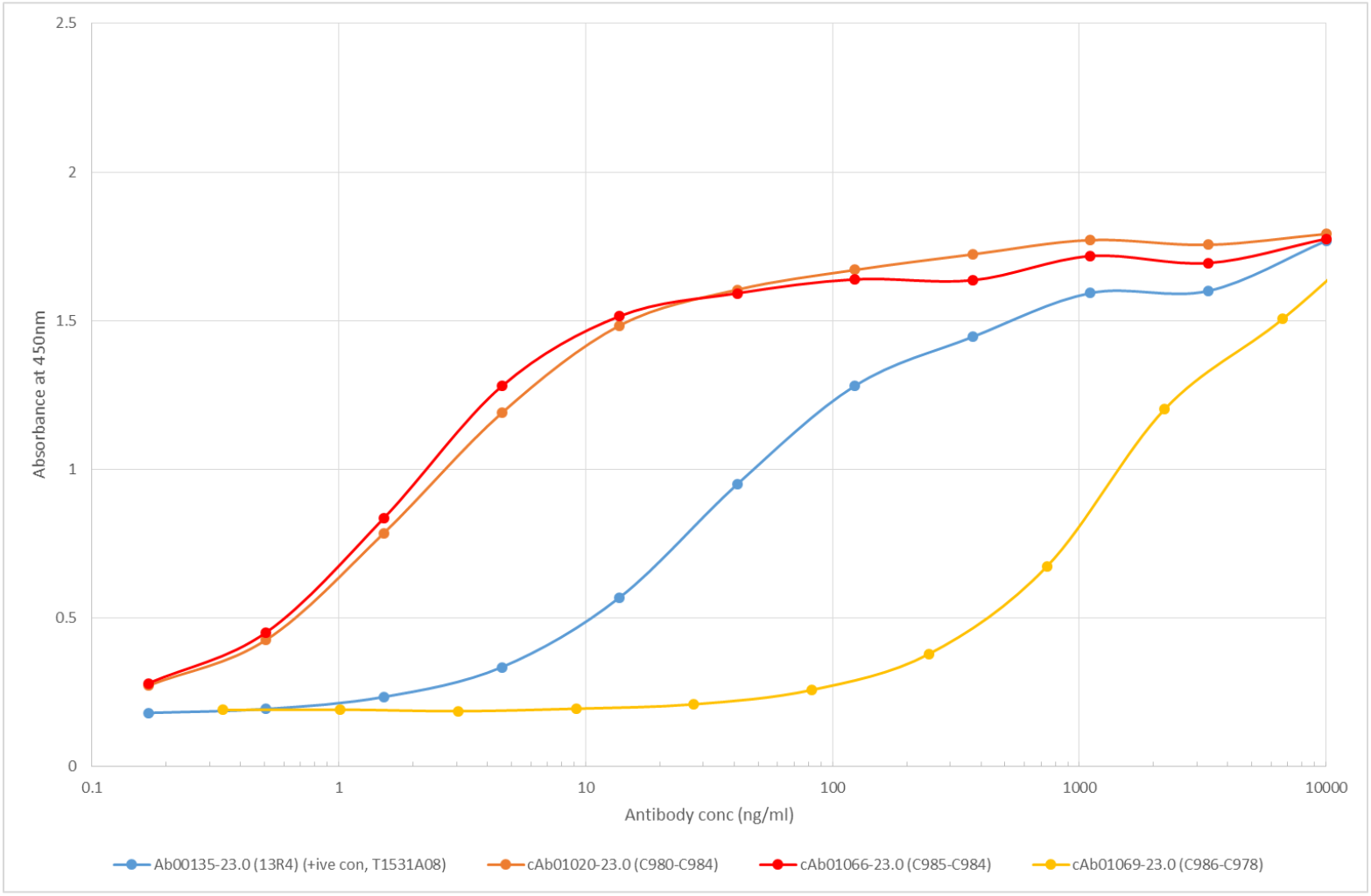
ELISA of monoclonal antibody candidates picked from recombinant Beta-gal fraction. Kd was calculated for c1020 and c1066, and Ab00135 is a Beta-gal positive control antibody. c1069 is an expressed candidate with little affinity for Beta-gal.

## H Hidden repertoire

Mutations are predicted and scored by aligning observed peptides to each unlabeled raw MS2 spectrum searching for single residue difference. Raw MS2 spectra are first deconvoluted using the approach from [24], and then converted to prefix-residue mass spectrum using predictive models trained on datasets from PXD002912 [30], PXD003868 [34], and PXD004948 [28] with procedures from [9]. Supposing that each MS2 spectrum represents a single peptide, a prefix-residue mass spectrum (prm spectrum) is defined as a list of mass and intensity pairs, where the intensities correlate with the accuracy that the corresponding masses represent peptide’s prefix masses.

To align a prm spectrum *S* to a peptide, the peptide is first converted to a theoretical prm spectrum *T*, and alignment is performed using a modified version of the Spectral Alignment Algorithm of Section 8.15 [17]. Briefly, prm pairs are matched along two diagonals, *D*_0_ and *D*_1_, such that any pair of prms on *D*_0_ have negligible mass difference and *D*_1_ have negligible difference from |*M_S_ − M_T_*|, the mass difference of total residue masses from both spectra. An optimal alignment is found by recursively calculating the maximum scoring residue path of prm pairs ending at *s, t* on either *D*_0_ or *D*_1_, where the *s′, t′* prior to *s, t* differ by residue masses. More formally, let *m*(*p*) be the mass of a prm, *R*(*p*) be the set of prms prior *p* in the spectrum that differ in mass by a mono- or di-residue mass, and *e* be an error tolerance term set to 0.02 Da. The optimal alignment is calculated as:

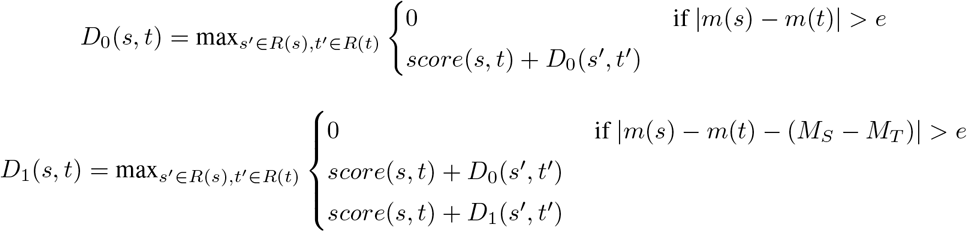

where

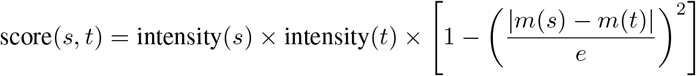

Adding terminal prm pairs, (0, 0) and (*M_S_, M_T_*) with *D*(0,0) = 1, allows for the optimal alignment score to be found recursively on *D*_1_(*M_S_, M_T_*). The algorithm runs in *O*(*rn*) where *n* is the number of prms in a spectrum and *r* is the most number of prms in *R*(*p*). A mutation is called by backtracking the alignment and finding the transition of (*s,t*) on *D*_1_ to (*s′, t′*) on *D*_0_, where the mutated residue has mass *s − s′*, and called on residue position *t*. As the spectral alignment does not account for richness of peaks in a spectrum, mutated peptide annotations were rescored with spectral probability, using MSGFDB generating score function [18].

To estimate precision and recall of mutation prediction and alignment, we simulate mutation finding by downsampling peptide spectrum matches from the database search results on affinity purified material from peptide and recombinant immunizations. In the downsampling, peptides are searched against prm spectra where some spectra represent mutated peptides. Briefly, a dataset is selected and peptides are randomly removed from the dataset if there exists more than one mutated peptide in the dataset. On average, the number of unique peptide pairs per simulation differing by a single residue is 390, 100, and 195 for recombinant Beta-gal, peptideA Beta-gal, and peptideB Beta-gal datasets. For each of these pairs, one peptide is assumed to be the mutated peptide, and a random spectrum matching the peptide is selected. The other peptide is kept, unmutated, to use for searching in the simulation. Remaining peptides are randomly partitioned into the spectra set and peptide sets. For the recombinant Beta-gal dataset, on average there are 390 mutated peptides out of 2600 spectra, and searched against 2600 peptides.

A spectral probability threshold of 10^−^8 results in average precision 0.66-0.69 and average recall 0.65-0.72 across the three datasets. Mutated peptide spectrum matches with < 10^−8^, are considered mutations. A number of false positives

**Figure S8:**
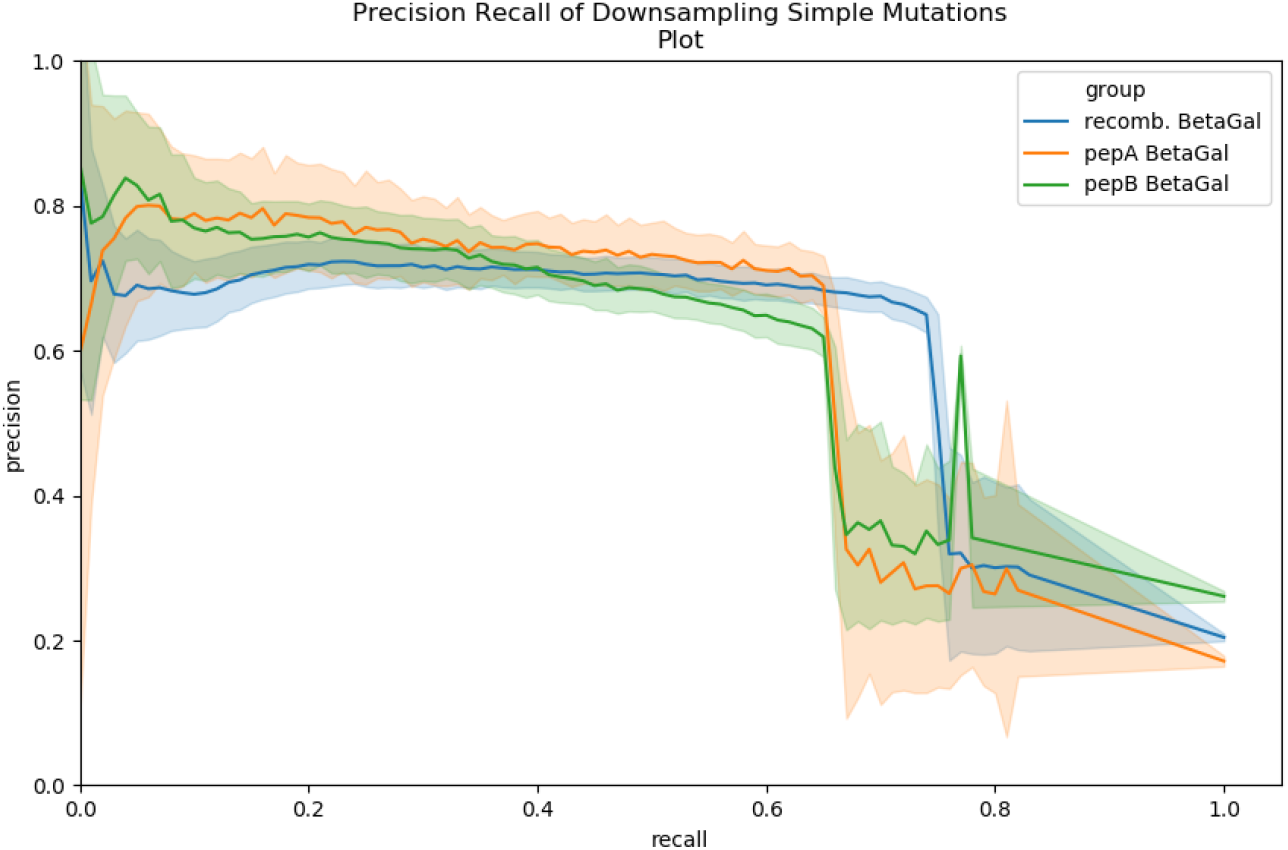
Precision recall curves fro downsampling of mutation pairs across 3 datasets, repeated 50 times, using the above algorithm to find single residue mutation and rescoring using MSGFDB spectral probability.

with low spectral probability are due to peptide pairs differing by more than one mutation. Supplemental Figure S9 shows that 70-90% of false positives mutated peptide calls have an underlying peptide pair with Levenshtein distance (edit distance) 2 to 4.

**Figure S9:**
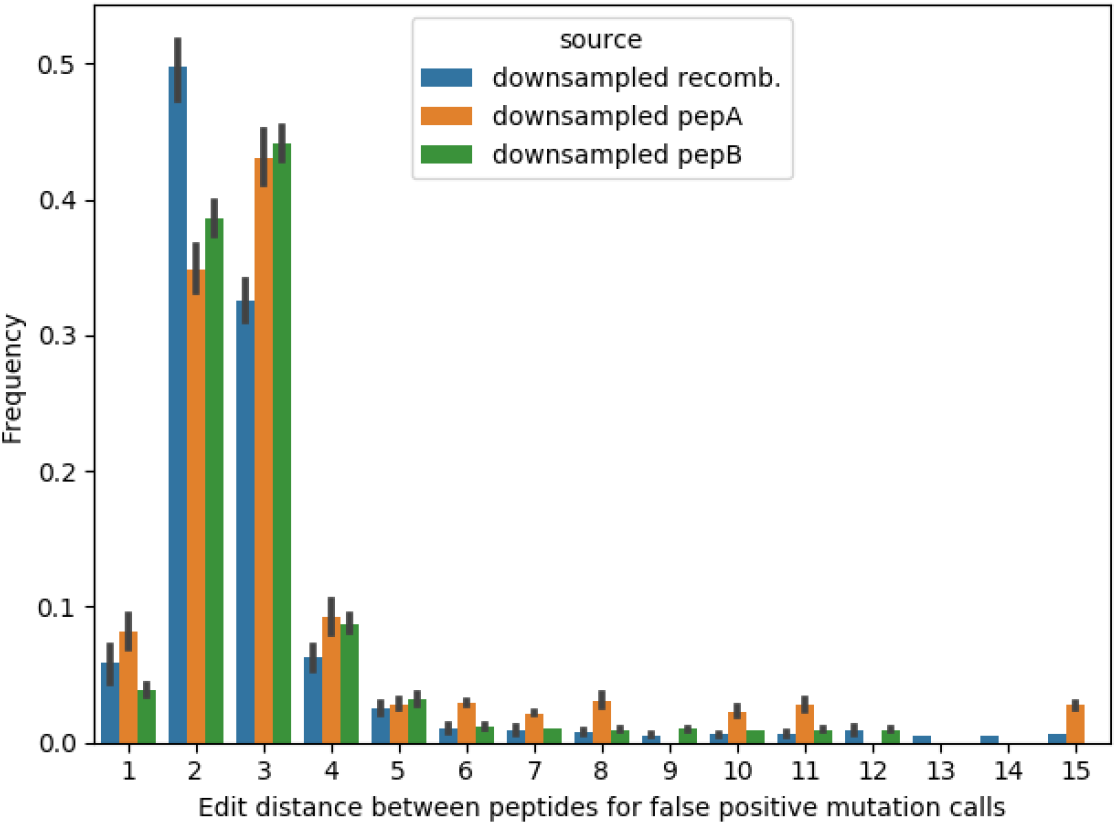
False positives explained by single mutation calls on peptides differing by 2-4 differences.

